# Substrate Curvature Influences Cytoskeletal Rearrangement and Modulates Macrophage Phenotype

**DOI:** 10.1101/2024.08.12.607593

**Authors:** Austin Sovar, Matthew Patrick, Ramkumar T. Annamalai

**Affiliations:** Department of Biomedical Engineering, University at Buffalo, Buffalo, NY 14260; Department of Biomedical Engineering, University of Kentucky, Lexington, KY 40506

**Keywords:** Microgel, Gelatin, Immunomodulation, Macrophages, Curvature

## Abstract

Inflammation serves as a critical defense mechanism against pathogens and tissue damage but can lead to chronic diseases, such as cardiovascular disease and diabetes, when dysregulated. Macrophages play a pivotal role in orchestrating inflammatory responses, transitioning from pro-inflammatory M1 to anti-inflammatory M2 phenotypes to resolve inflammation and promote tissue repair. Current approaches to modulate macrophage phenotype predominantly rely on biochemical cues, which may induce systemic side effects. Given the mechanosensitivity of macrophages, this study investigates biophysical cues, specifically substrate curvature, as a localized strategy to regulate macrophage phenotype and minimize systemic repercussions.

We hypothesized that substrate curvature influences macrophage immunophenotype by modulating F-actin polymerization. To test this hypothesis, we fabricated spherical microgels with tunable curvatures and characterized their biophysical properties. Our findings indicate that macrophages adhere to microgel surfaces irrespective of curvature, but the curvature significantly alters F-actin dynamics. Furthermore, manipulating cytoskeletal dynamics via selective actin inhibition partially reversed curvature-induced changes in macrophage phenotype. These results underscore the pivotal role of substrate curvature in modulating macrophage behavior and immunophenotype.

Overall, our study demonstrates that substrate curvature significantly influences macrophage cytoskeletal dynamics and resulting immunophenotype. This simple approach can be utilized as a localized immunomodulatory treatment for inflammatory diseases.

## 1. Introduction

Inflammation is a critical component of the immune system, serving as the body’s first line of defense against foreign threats or injurious stimuli. It plays a vital role in fighting infections, clearing damaged cells, and initiating tissue repair. When effectively regulated, the inflammatory response resolves threats promptly and transitions into a healing state. However, when dysregulated, the inflammation can become prolonged, impeding repair, exacerbating tissue damage and promoting fibrosis. This chronic inflammatory response is a fundamental aspect of several degenerative and chronic conditions, including diabetes, cardiovascular disease, arthritis, and various organ-specific diseases^1-3^ that affect populations globally. The World Health Organization (WHO) recognizes chronic inflammation-related diseases as a significant threat to global health. For instance, in the US alone, cardiovascular disease accounts for 800,000 deaths annually, and around 30.3 million people suffer from diabetes^1^. These conditions often share a common underlying mechanism involving dysregulated inflammatory and immune responses, highlighting the urgent clinical need for therapies to mitigate these harmful processes.

The initiation, progression, and resolution of the immune response are heavily dependent on the innate immune cells, particularly macrophages, which play a central role in modulating inflammation^4^. Macrophages are highly plastic cells that respond to the local microenvironment by propagating signals essential to each stage of the immune response through phenotypic changes. These phenotypes range from pro-inflammatory (M1) to anti-inflammatory (M2), with a spectrum of specialized responses in between^5-7^. Although It is well-known that macrophage polarization can be influenced by both biochemical and biophysical cues, the precise mechanisms underlying these effects remains poorly understood^8^. A deeper understanding of these mechanisms would enable the harnessing of macrophage plasticity to modulate their immunophenotype, potentially resolving dysregulated immune responses.

Numerous studies have demonstrated successful approaches to modulating macrophage immunophenotypes^9-13^. Systemic application of biochemical agents, such as JAK inhibitors, which inhibit the activity of Janus kinases^14^ and proinflammatory cytokines such as IL-1β and TNFα^15, 16^, have shown efficiency in alleviating chronic inflammation. Similarly, humanized monoclonal antibodies targeting IL-6R to antagonize IL-6 binding and hinder inflammatory responses are currently utilized in treating several inflammatory conditions^17^. However, these biochemical factors often lack target specificity and can increase susceptibility to infection and systemic toxicity^18^. This underscores the need for targeted and localized approaches that can modulate the immune response without relying on soluble factors.

Biophysical cues such as substrate stiffness^19^, viscoelasticity^20^, geometry^21^, and spatial patterns^22^ have shown the potential to address this challenge^23^. Cells primarily recognize these physical cues through integrin-mediated mechanotransduction pathways, which convert mechanical forces into biochemical signals, subsequently altering cytoskeletal dynamics and phenotype^24^. Macrophages, in particular, respond to these physical cues through actin-cytoskeletal reorganization, nuclear deformation, and subsequent gene expression (**Fig.1**)^25^.

**Figure 1.**
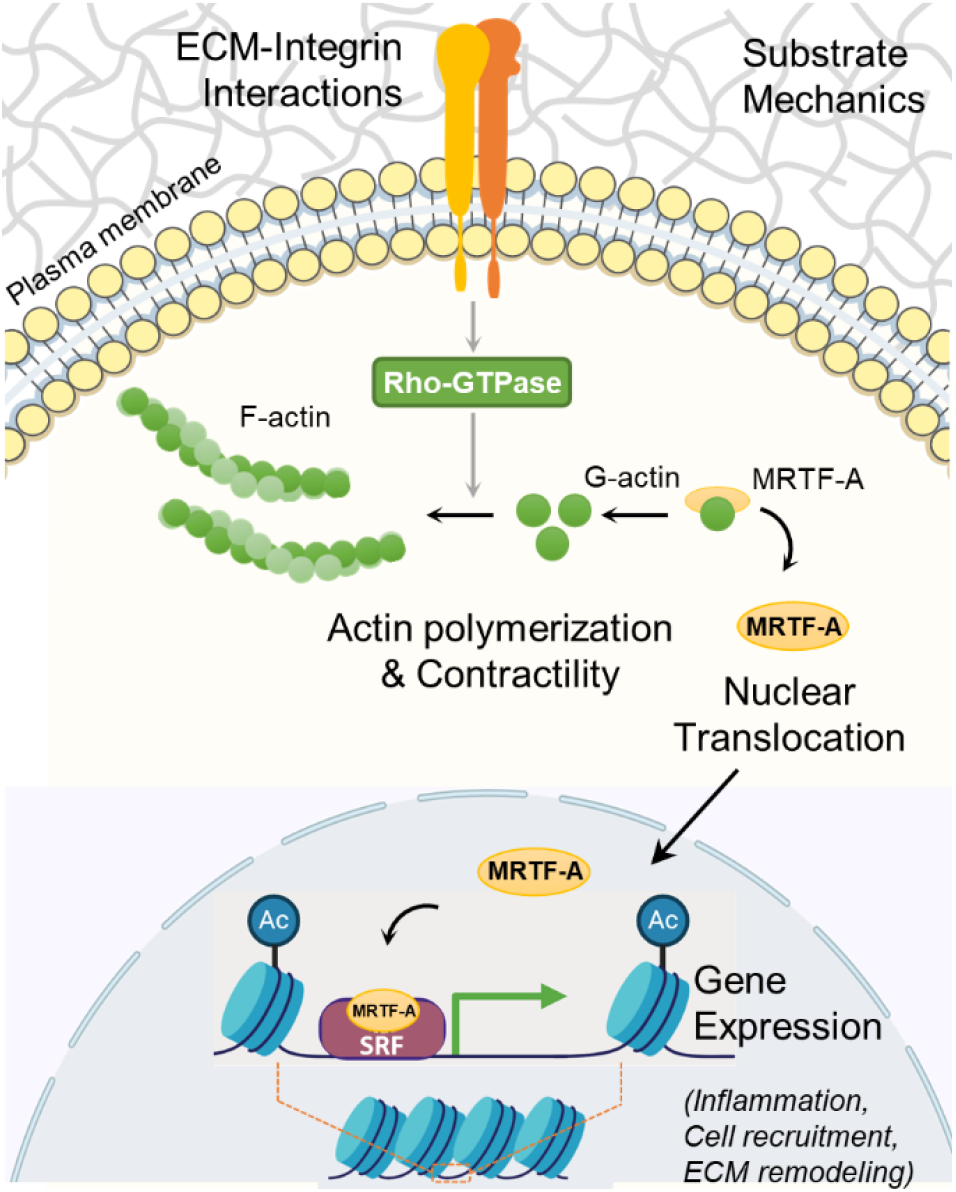
Cytoskeletal Dynamics Influences Immunophenotype of Macrophages. Physical forces in the form of substrate mechanics elicit transcriptional control of macrophage phenotype and function through F-actin polymerization and subsequent nuclear translocation of transcription factors.

Here, we describe a straightforward approach to modulate macrophage responses using simple gelatin-based microgels with tunable properties, including size and curvatures, capable of inducing a localized immunomodulatory effect. We hypothesize that the substrate curvature can modulate macrophage immunophenotype through F-actin polymerization. To test the hypothesis, we fabricated tunable spherical microgels with varying curvatures and characterized their biophysical properties. Our findings indicate that macrophages readily attach to the surface of these microgels regardless of their curvature. However, the curvature of the substrate significantly influences F-actin polymerization, leading to changes in macrophage phenotype. This effect was partially reversed when treated with selective actin inhibitor Latrunculin A. These results demonstrate the influence of substrate curvature on macrophage cytoskeletal dynamics and resulting immunophenotype. Consequently, this simple approach has the potential to be utilized as a localized immunomodulatory treatment for inflammatory diseases.

## 2. Methods

### 2.1. Microgel fabrication

Microgels were fabricated as previously described^26^, Briefly, a 6% gelatin (type A, 300 bloom, Sigma) in deionized (DI) water stock solution was created. This solution was dispensed dropwise into a polydimethylsiloxane (PDMS, viscosity = 100 cS, Clearco Products Co., Inc.) bath heated to 37° C. The mixture was emulsified with a double impeller at either 500 or 1000 rpm to achieve the desired size range. After 5 minutes of emulsification at 37° C, the mixture was cooled using an ice bath and stirred for 30 minutes. The microgels were then separated through centrifuging at 185 g for 5 minutes. The supernatant was discarded, and the resulting microgel pellet was washed thrice with Dulbecco phosphate buffer solution (PBS, Invitrogen) supplemented with 1% TWEEN 20 (Sigma). The microgels were then crosslinked with 1 wt% genipin in PBS for 6 or 48 hours to achieve different crosslinking densities. After cross-linking, the excess genipin was removed by washing with 100% ethanol and stored in 100% ethanol at 4-8° C until further use.

To obtain microgels of specified size and curvature, they were washed and swollen in PBS, sonicated for 5 minutes in an ice bath, and sorted using nylon mesh filters. For our studies, 40-50 µm (κ = 0.045 µm-1), 150-250 μm, and 350-400 μm (κ = 0.005 µm^-1^) nylon meshes were used. This procedure was repeated twice to remove aggregates thoroughly.

### 2.2. Swelling and polymer density

Microgel swelling and polymer density were characterized using microgels within the size range of 150-250 μm crosslinked for 6 hours (low crosslinking) and 48 hours (high crosslinking). The mass and volume swelling ratio were determined and used to find the polymer densities. The swelling ratios were calculated through quantitative measurements from hydrated and dehydrated microgels. For hydration, the microgels were swollen in 10 mM PBS before measurements. Dehydration was achieved through flash freezing in liquid nitrogen followed by lyophilization. The polymer density was calculated using the following equation:

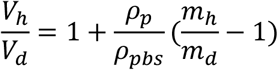

Where *V*_*h*_ is the volume of hydrated spheres, *V*_*d*_ is the volume of dry spheres, *ρ*_*p*_ is the polymer density, *ρ*_*pbs*_ is the density of microgels after swollen in PBS, *m*_*h*_ is the mass of hydrated spheres, and *m*_*d*_ is the mass of the dehydrated spheres. Bright field images of the microgels were obtained and processed in ImageJ to measure the diameter and calculate the volume of spheres.

### 2.3. Compression testing

The mechanical properties of the microgel formulations were characterized and compared using compression testing. Briefly, cylindrical bulk gels were prepared in a 24-well flat bottom plate, crosslinked with 1 wt% genipin in PBS, washed with ethanol at specific time points to stop the reactions, and then rehydrated in 10 mM PBS for testing. Uniform disks of 5 mm in height were cut out using a 6 mm biopsy punch. The cylindrical samples were subjected to compressed testing using a Universal Testing Machine (Instron). The samples were compressed at a strain rate of 10 mm/min, generating a force-displacement curve. The elastic modulus was then calculated from the linear region (between 5-15% strain region) of the resulting stress-strain curve.

### 2.4. Cell culture

A murine macrophage cell line IC-21 (ATCC) was cultured with RPMI-1640 Medium (ATCC) supplemented with 10% fetal bovine serum, anti-biotic and anti-mycotic supplements. Cells were expanded and maintained at 37°C in a CO_2_ incubator. Macrophages were grown to ∼80% confluency then detached and used for experimental use.

For curvature studies, microgels (∼3 mg/mL) of the desired curvature (40-50 µm for low curvature and 350-400 µm for high curvature) were seeded with ∼3 million macrophages. To prepare the microgels for cell seeding, they were swollen with media in vented culture tubes (VWR), which were then replaced with macrophages suspended in growth media. The seeded microgels were kept in the incubator and gently mixed every 10 minutes for 1 hour to prevent aggregation. They were then washed with fresh media to remove unattached cells and transferred to a 35 mm bioinert µ-dishes (Ibidi) to prevent aggregation. For Latrunculin A (LatA) studies 100 nM of LatA was added to samples 24 hours before harvesting samples.

### 2.5. RNA Isolation and Quantitative Gene Expression

Total RNA was isolated from samples by first removing media from the suspension and adding 500 µL of TRIzol. The mixture was then vortexed vigorously, centrifuged at 10,000 g for 5 minutes, and the supernatant was transferred to a new 1.5 mL tube. Subsequently, 100 µL of chloroform was added, mixed vigorously, incubated at room temperature for 10 minutes, and then centrifuged at 16,000 g at 4°C for 10 minutes. The aqueous phase was transferred to a new 1.5 mL tube where 10 µg of glycogen and 250 µL of isopropanol were added. This mixture was then mixed, incubated at room temperature, and centrifuged at 12,000 g at 4°C for 10 minutes. The supernatant was discarded, and 500 mL of 75% ethanol was added to the remaining pellet, which was then vortexed and centrifuged at 7,500 g at 4°C for 10 minutes. After discarding the supernatant, the pellet was allowed to air dry. 10-20 µL of RNAse-free water was added to the dry pellet, and the solution was heated at 55°C for 15 minutes. The quality of isolated RNA was assessed through absorbance measurement using NanoDrop (ThermoFisher).

With isolated RNA samples, qPCR gene expression analysis was conducted. In each qPCR well, RNA samples were combined with RNAse free water, 2x reaction buffer, ROX, SuperScript, and TaqMan probe in a 10:50:2:2:31:5 ratio, respectively. Various TaqMan probes were employed to identify distinct gene expressions. The Ct values of *Gapdh* (Mm99999915_g1), *Actb* (Mm02619580_g1), and *Hsp90ab1* (Mm00833431_g1) were averaged and used as internal controls. *Arg1* (Mm00475988_m1) and *Igf1* (Mm00439560_m1) were used to characterize M2-prohealing genes. *Il-1b* (Mm00434228_m1), *Nos2* (Mm00440502_m1), and *Tnf* (Mm00443260_g1) were used to characterize M1-proinflammatory genes. Arpc2 (Mm01254383_m1), Cfl1 (Mm03057591_g1), and Tln1 (Mm00456997_m1) were used to quantify changes in cytoskeletal proteins expressions. Expression of genes seeded on high and low curvature microgels for 1 day were used as the baseline for each respective group. Gene expression analysis was performed using QuantStudio 3 real-time PCR system (ThermoFisher).

### 2.6. Electron Microscopy

Samples intended for electron microscopy were fixed with glutaraldehyde. The fixed samples were dehydrated in 100% ethanol, flash-frozen in liquid nitrogen, and lyophilized. The Leica ACE 600 was used to sputter coat the lyophilized samples with a 5 nm-thick layer of platinum. Images were taken using the FEI Quanta 250 environmental scanning electron microscope (SEM).

### 2.7. Immunofluorescent Staining and Confocal Imaging

Samples intended for immunofluorescent staining were fixed with Z-fix (Anatech). Prior to staining, fixed samples were permeabilized with 0.1% Triton X in PBS for 3-5 minutes and then washed with 10 mM PBS. The samples were stained with Phalloidin-FITC (Molecular Probes) and DAPI (Molecular Probes) to target the F-actin cytoskeleton and DNA in the nucleus, respectively. The staining solution consisted of Phalloidin at 1:40 and DAPI at 1:1000 ratio with 1% BSA in PBS. Samples were stained for 30 minutes in the dark and washed twice with PBS.

Z-stack confocal images were acquired using a Nikon A1R confocal microscope and processed using ImageJ software. F-actin density was quantified by demarking macrophage boundaries and measuring the integrated fluorescence density (IntDen = sum of the grey scale values of each pixel * area of one pixel) expressed in arbitrary fluorescence units (AU). For cell clusters where it was challenging to demarcate the individual cell borders, the total F-actin density of the cluster was measured and averaged over the number of DAPI-stained nuclei in the cluster. All samples were imaged using consistent confocal camera settings for comparison.

### 2.8. Statistics

Data for this experiment were completed in triplicate, at least. Finalized measurements were analyzed as averages with standard deviations to represent error. A two-tailed, equal variance Student’s t-test was used to analyze single statistical comparisons. For multiple comparisons a two-way ANOVA was performed with a subsequent Tukey test. The Pearson correlation coefficient (r) was used to evaluate the linear correlation between two variables and the t-distribution to evaluate the associated p-value. For all tests, a p-value of less than 0.05 was considered significant.

## 3. Results

### 3.1. Tunable Microgels and their Physical Properties

Microgels were fabricated using a simple water-in-oil emulsification of solubilized gelatin followed by genipin crosslinking to the desirable extent. The size distribution of the microgels varied with the set speed of the propeller during the emulsification. Lower speeds (<500 rpm) generally resulted in microgels of a larger size range (150-400 µm), while higher speeds (>1000 rpm) resulted in smaller microgels (40-50 µm). To achieve narrow size ranges, nylon meshes were used to sort microgels to desired size ranges (**Fig.2A**, Hydrated 150-250 µm microgels). To achieve at least one-order-of-magnitude separation between the surface curvatures, we created microgels with two size ranges: 40-50 µm (κ = 0.045 µm-1) and 350-400 μm (κ = 0.005 µm^-1^). A mid-size range of 150-250 μm for material characterization purposes.

**Figure 2.**
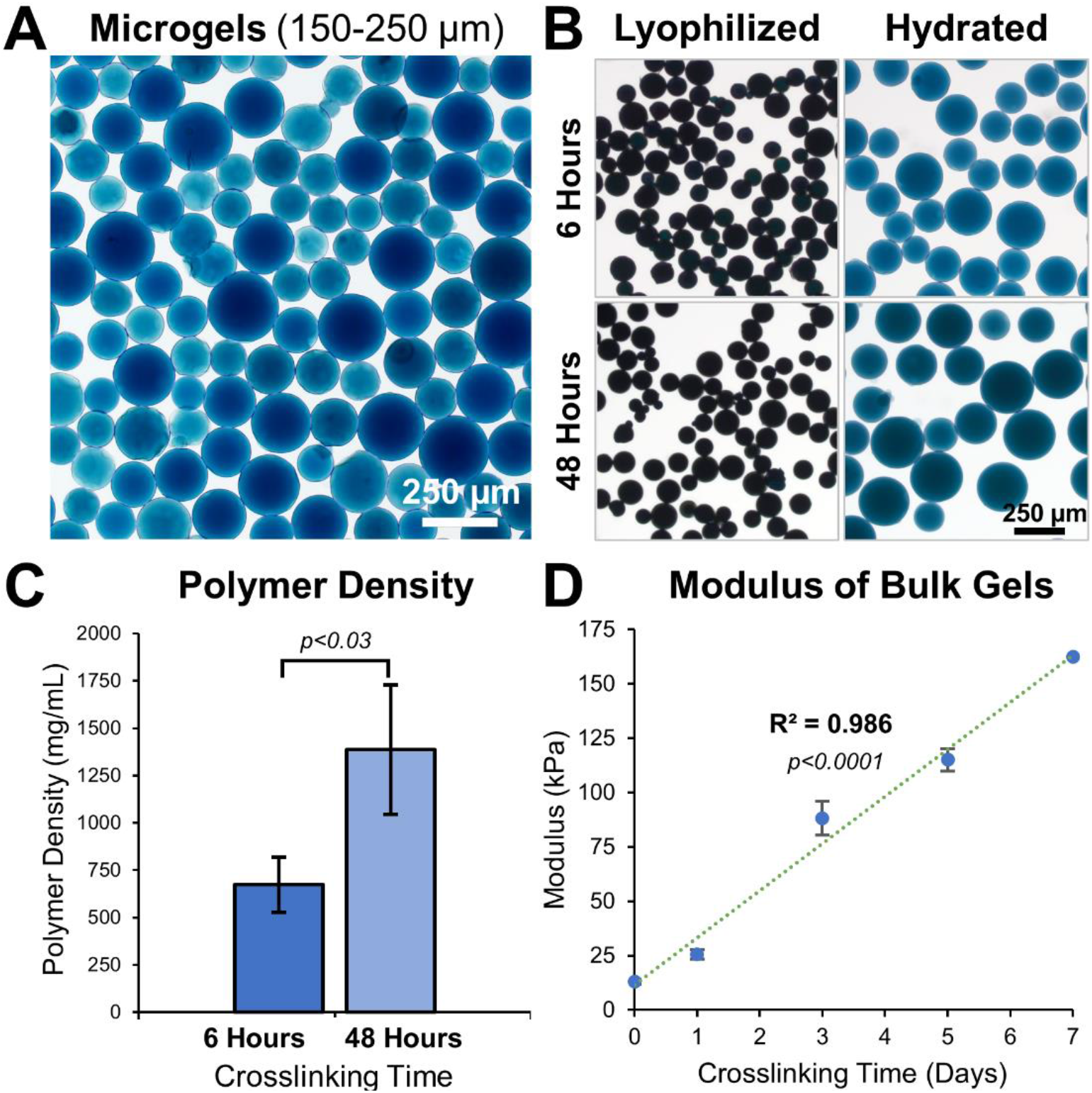
Tunable Microgel Fabrication and Characterization. A) Bright-field images of 48 hour crosslinked hydrated microgels of size range 150-250 μm. B) Images of 6 hours and 48 hours crosslinked microgels before and after hydration in PBS. C) Polymer density of 6 hours and 48 hours crosslinked microgels, n = 3 batches. D) Young’s modulus of gelatin hydrogel disks show a linear increase with crosslinking time, n = 6.

The material properties of a hydrogel can considerably influence cell behavior. By adjusting the crosslinking time, the microgel properties were easily fine-tuned (**Fig.2B**). Microgels crosslinked for 6 hours had a lower polymer density (674 ± 146 mg/mL) than microgels crosslinked for 48 hours (1386 ± 342 mg/mL, **Fig.2C)**. Microgels with low crosslinking (6 hours) had a smaller volume-swelling ratio of 502 ± 68.7 % and microgels with high crosslinking (48 hours) had a larger volume-swelling ratio of 855 ± 82.2% (**Fig.2B**).

Since the microgels were too small for mechanical testing, we used bulk hydrogel samples crosslinked for different periods and performed compression testing to study their bulk properties. Solidified gelatin hydrogel disks, 5 mm high and 6 mm in diameter, were crosslinked for varying periods and subjected to compression tests. The elastic modulus of the disks ranged between 13.0 ± 1.22 KPa at day 0 and 163 ± 5.29 KPa at day 7 of crosslinking time, exhibiting a linear correlation (**Fig.2D**, R^2^ = 0.986, p<0.0001, n=6).

### 3.2. Substrate Curvature Moderately Influences Macrophages Attachment and Spreading

To investigate the effect of substrate curvature on macrophages cytoskeletal dynamics and phenotype, microgels with two degrees of curvature were selected: high curvature microgels with a diameter of 40-50 µm (κ = 0.045 µm^-1^, **Fig.3A(i)**) and lower curvature microgels with a diameter of 350-400 µm (κ = 0.005 µm^-1^, **Fig.3A(ii)**). These microgels were crosslinked for 48 hours to maintain a consistent modulus of around 50 kPa. After thorough washing with ethanol, followed by PBS and culture media, the microgels were seeded with macrophages in suspension cultures.

**Figure 3.**
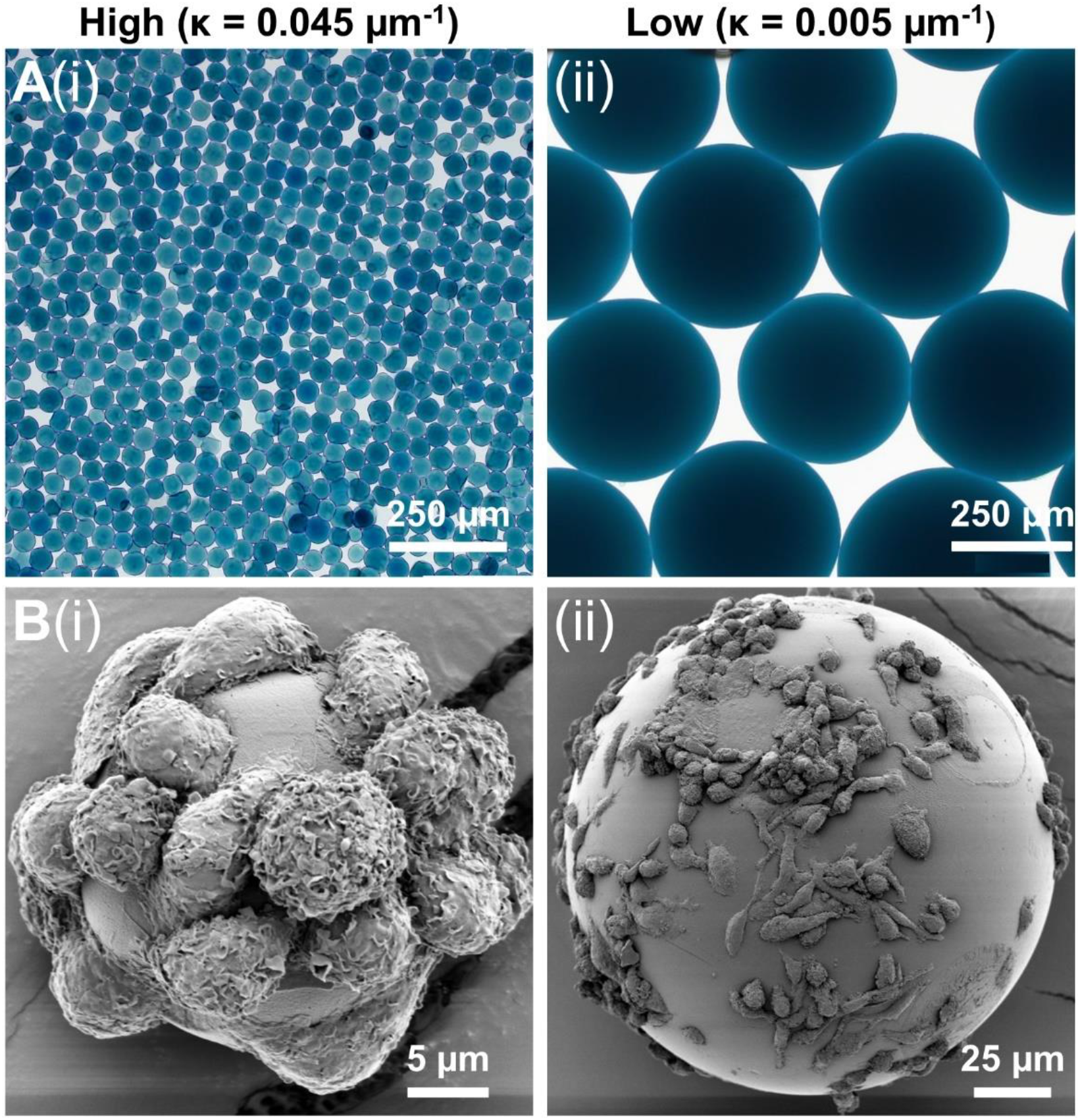
Macrophage Attachment on Low and High Curvature Microgels. **A)** Bright-field microscopy images of the (i) high curvature (κ = 0.045 µm^-1^) and (ii) low (κ = 0.005 µm^-1^) microgels showing uniform smooth surface morphology, **B)** SEM images of macrophages seeded on (i) high curvature and (ii) low microgels showing morphological differences.

The macrophages quickly attached to the gelatin surface, covering both high and low curvature microgels within 24 hours. No significant difference in cell attachment was noticed between the high (**Fig.3B(i)**) and low (**Fig.3B(ii)**) curvature conditions. But the attached macrophages tended to be more elongated in the low curvature compared to the high curvature microgels (**Fig.3B(ii)**). On the high curvature microgels, macrophages tended to warp around the microgel, displaying a more compact morphology.

### 3.3. Substrate Curvature Modulate Cytoskeletal Dynamics and Phenotype of Macrophages

To investigate the influence of substrate curvature on cytoskeletal dynamics and corresponding phenotypic changes, macrophages were cultured on low and high curvature microgels for 7 days and characterized through actin cytoskeleton staining and gene expression studies. Fluorescence staining and quantification of the cytoskeletal F-actin, on day 7, revealed that the macrophages on low curvature microgels exhibited a 64% higher F-actin density (898.87 ± 128.36 AU, **Fig.4A(i),C**) compared to those on the high curvature microgels (548.70 ± 127.23 AU, *p<0*.*0001*, n=25, **Fig.4A(ii),C**).

**Figure 4.**
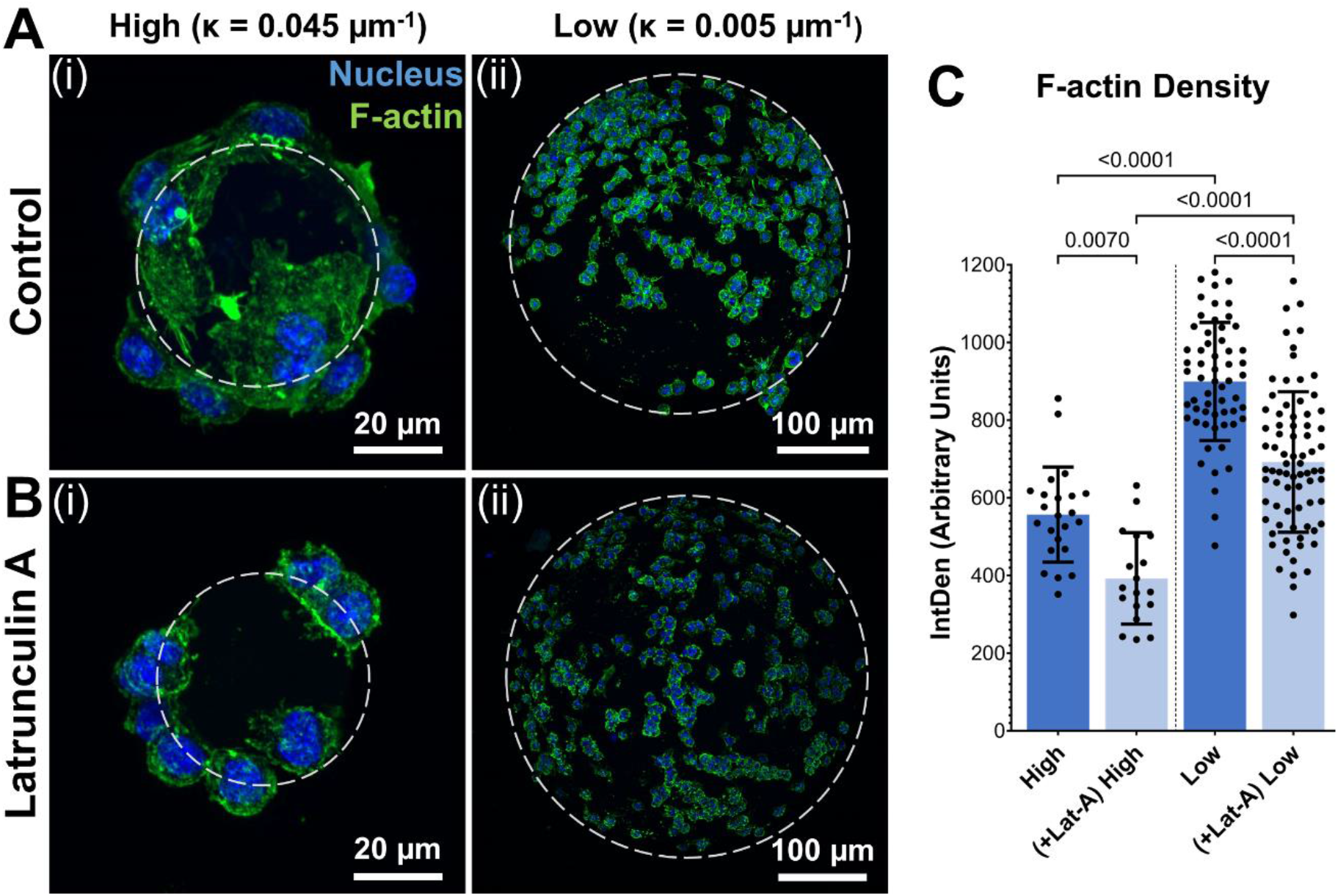
Substrate Curvature Modulate Cytoskeletal Dynamics. **A)** Confocal stacks of Day 7 macrophages seeded on high (i) and low (ii) curvature microgels. **B)** Confocal stacks of macrophages seeded on high (i) and low (ii) curvature microgels after latrunculin-A treatment for the last 24 hours of day-7 cultures. Green and blue fluorescence corresponds to cytoskeletal F-actin and nuclei, respectively. **C**) F-actin density (IntDen) in macrophages seeded on microgels in the respective groups (n = 25) measured through ImageJ quantification (n>25 for each group).

For further validation, a selective inhibition of actin polymerization was performed using latrunculin-A (Lat-A) treatment for the last 24 hours of the 7-day cultures. Lat-A treatment significantly reduction F-actin density in both groups (F-actin density in Low = 725.12 ± 148.12 AU, *p<0*.*0001* and High = 404.69 ± 113.7 AU, *p<0*.*0001*) and led to a corresponding reduction in cell spreading in both groups (**Fig.4B, C**). Despite this reduction, the low curvature microgels maintained significantly (79%) higher F-actin density compared to high curvature microgels, indicating a strong influence on the cytoskeleton of macrophages.

Subsequently, we investigated the changes in immunophenotype of macrophages caused by alterations in actin polymerization. Actin polymerization is known to trigger the release of the transcription factor MRTF-A^24^, which regulates expression levels of several genes involved in inflammatory responses. A panel of prominent inflammatory and anti-inflammatory genes, as well as genes implicated in actin cytoskeletal dynamics, was identified. The expression levels of these genes were quantified and compared after 1 week of culturing in low and high curvature microgels.

The relative expression levels of several key genes exhibited significant differences between macrophages cultures on low and high curvature microgels at day 7, both before and after Lat-A treatment (**Fig.5A**). Notably the pro-inflammatory gene *Tnf*, and the anti-inflammatory genes *Arg1* and *Nos2* showed robust curvature dependent expression patterns.

**Figure 5.**
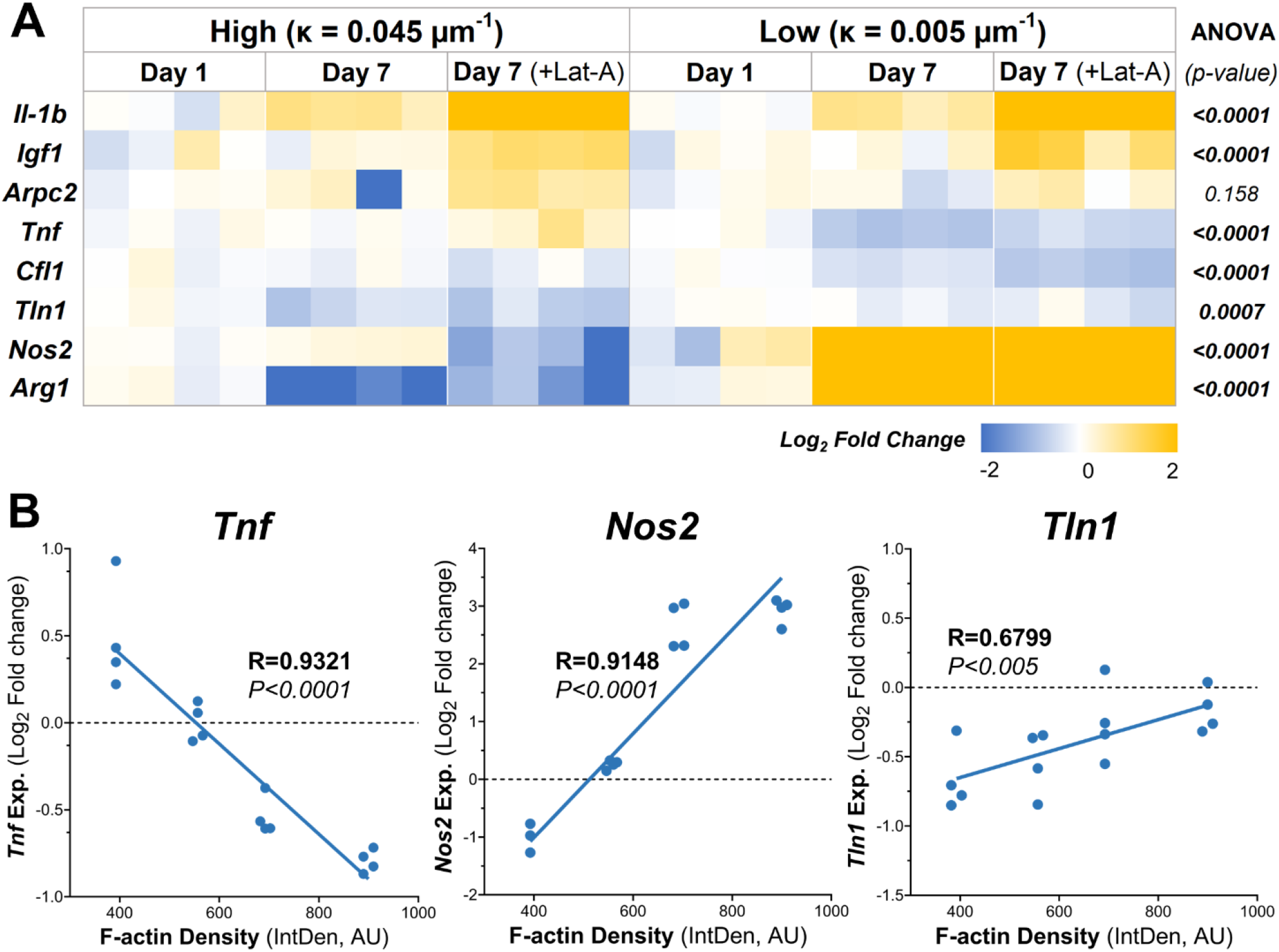
Substrate Curvature Modulate Macrophage Phenotype. **A)** The relative gene expression levels of common inflammatory and anti-inflammatory markers in macrophages cultured on high and low curvature gels for 7 days, treated and untreated with Lat-A (n = 4). **B**) Correlation plots of gene expression at day 7 with significant linear relationships with f-actin level. AU-Arbitrary units.

Low curvature condition led to a significant downregulation of *Tnf* (1.7-fold decrease, *p=0*.*0001*). Conversely, *Arg1* was significantly downregulated in high curvature conditions (4-fold, *p=0*.*0103*), while *Arg1 and Nos2 was* being significantly upregulated in low curvature conditions (7.7-fold, *p=0*.*0003*, and 7.6-fold *p<0*.*0001*, respectively). Interestingly, these trends persisted even after Lat-A treatment that significantly reduced the F-actin polymerization. Correlation analysis revealed a robust and significant linear correlation between F-actin levels and the expression of *Tnf* (R=0.9321, *p<0*.*0001*), *Arg1* (R=0.9148, *p<0*.*0001*), and *Tln1* (R=0.6799, *p<0*.*005*) (**Fig.5B**), indication a strong link between cytoskeletal dynamics and macrophage phenotype.

**Table 1.** Pairwise comparison of treatment conditions. (p<0.05)

| **Gene** | **High Curvature** |  |  | **Low Curvature** |  |  | **High vs Low** |  |
| --- | --- | --- | --- | --- | --- | --- | --- | --- |
|  | Day 1 vs -Day 7 | Day 1 vs Day 7 (+Lat-A) | Day 7 vs Day 7 (+Lat-A) | Day 1 vs -Day 7 | Day 1 vs Day 7 (+Lat-A) | Day 7 vs Day 7 (+Lat-A) | Day 7 | Day 7 (+Lat-A) |
| ***Il1b*** | **0.0006** | **<0.0001** | **<0.0001** | **0.001** | **<0.0001** | **<0.0001** | 0.9999 | **<0.0001** |
| ***Igf1*** | 0.995 | **0.0033** | **0.0099** | 0.9908 | **0.0014** | **0.0048** | >0.9999 | 0.9982 |
| ***Arpc2*** | 0.9729 | 0.5282 | 0.1307 | 0.9997 | 0.9327 | 0.8221 | 0.9946 | 0.9292 |
| ***Tnf*** | >0.9999 | **0.0087** | **0.0088** | **<0.0001** | **0.0032** | 0.3088 | **<0.0001** | **<0.0001** |
| ***Cfl1*** | 0.9774 | 0.2664 | 0.6521 | **0.0206** | **<0.0001** | **0.0171** | 0.0857 | **0.0009** |
| ***Tln1*** | **0.0178** | **0.0028** | 0.9473 | 0.8574 | 0.5162 | 0.9887 | 0.1634 | 0.1018 |
| ***Nos2*** | 0.9763 | **0.0103** | **0.0022** | **<0.0001** | **<0.0001** | 0.9728 | **<0.0001** | **<0.0001** |
| ***Arg1*** | **<0.0001** | **0.0003** | 0.1491 | **<0.0001** | **<0.0001** | **0.0409** | **<0.0001** | **<0.0001** |

## 4. Discussion

Inflammation is a critical and tightly regulated aspect of the immune response, and its dysregulation often leads to debilitating diseases^1-3^. While small molecules, such as NSAIDs^18^, have effectiveness in modulating inflammatory responses, they can cause significant off-target effects, such as gastrointestinal complications^27^ and systemic issues. This underscores the need for alternative therapeutic approaches that do not rely solely on biochemical interventions.

It is well recognized that mechanical forces play a vital role in regulating major cellular events, including inflammatory response, tissue development, regeneration, and healing after trauma^28^. However, majority of the work has focused on inflammation mediated by biochemical signals, with less attention given to the physical parameters of the inflammatory environment, especially matrix mechanics. These physical parameters, particularly matrix stiffness and curvature, are crucial in modulating immunophenotypes and potentially key to developing the next generation of therapies.

In this study, we address this gap by investigating how substrate curvature influence macrophage behavior and immunophenotype. Macrophages are innate immune cells known for their significant influence on the inflammatory response. They exhibit remarkable plasticity in response to both biochemical and biophysical cues. This plasticity allows them to both initiate and resolve inflammation, making them ideal candidates for therapeutic manipulation. By elucidating the impact of physical properties like curvature on macrophage activation, we open new avenues for therapeutic interventions that minimize off-target effects and harness the potential of biophysical cues to regulate inflammation. This approach could lead to more precise and effective treatments for inflammatory diseases.

### 4.1. A simple and tunable microgel system

To study the effect of curvature, we used the well-characterized biomaterial gelatin, crosslinked with biocompatible genipin, to create microspheres of various sizes. Gelatin was chosen as the base for our microgel system due to its well-characterized biocompatibility and naturally occurring integrin binding sites, which facilitate cell attachment without the need for chemical modification. Our characterization of the microgels revealed that both stiffness and polymer density can be precisely tuned by altering the crosslinking time. Given that these material properties can significantly influence cell behavior^30, 31^, it is crucial to evaluate and maintain their consistency. Therefore, all microgels were crosslinked for 48 hours to achieve near-complete crosslinking, ensuring minimal variation between experimental conditions. Additionally, the curvature range of the microgels can be easily set through sorting. Previously, we showed the potential of the microgels for both experimental investigation and therapeutic use^32, 33^. They are fully degradable, allowing them to integrate with the native ECM while maintaining their original structure for a few weeks. These microgels enable high-throughput analysis of a wide range of material characteristics while possessing valuable properties for an effective therapeutic treatment option.

### 4.2. Curvature-driven actin cytoskeletal dynamics

The integrin-rich surface of the microgels facilitated rapid macrophage adhesion. Once attached, the macrophages exhibited distinct curvature-dependent morphology and behavior. This morphological difference is likely due to the varying bending energy required for cells to adhere to surfaces of different curvatures. Typically, cells avoid migrating or adhering to high curvature surfaces due to the increased energy required to wrap around and attach. However, spheres compel cells to attach and expend the necessary bending energy due to their uniform curvature^34^. Similar studies have shown that cell spreading is enhanced in lower curvature surfaces and restricted on higher curvature ones, consistent with our observations^35^. The low-curvature microgels required less bending energy, allowing macrophages to spread more efficiently, elevating F-actin polymerization.

### 4.3. F-actin polymerization and transcriptomic modulation in macrophages

F-actin polymerization requires monomeric G-actin, which unbinds from myocardin-related transcription factor A (MRTF-A) to become active^36^. The unbound MRTF-A then translocates to the nucleus, binds with serum response factor (SRF), and activates the transcription of specific genes, including inflammatory genes *Tnf, Nos2*, and *Il6*^37^. We hypothesized that this actin-dependent gene expression influences macrophage phenotype. Thus, we compared macrophages cultured on both high- and low-curvature microgels and evaluated changes in inflammatory and cytoskeletal gene expression in response to differential F-actin polymerization.

Our results showed that macrophages on low-curvature microgels exhibited higher F-actin levels compared to those on high-curvature microgels after one week of culturing. Latrunculin A (LatA), that inhibits F-actin polymerization, proportionally reduced F-actin levels in both groups. This allowed us to assess the influence of F-actin on macrophage phenotype at differential F-actin levels and presumably the subsequent MRTF-A nuclear translocation as well. Two inflammatory genes exhibited strong linear relationships with F-actin levels: *Tnfα* showed a strong negative correlation, while *Nos2* showed a strong positive correlation. *Tnfα* is a major component of the innate inflammatory response, initiating the inflammatory cascade by promoting the expression of multiple pro-inflammatory proteins^38^. *Nos2* exhibits a pro-inflammatory effect at low concentrations, while at higher concentrations can mount an anti-inflammatory effect by promoting the apoptosis of inflammatory cells^39^. Additionally, the cytoskeletal gene, *Tln1* (talin-1), essential for linking activated integrins to cytoskeletal actin and focal adhesion assembly^40^, shows a strong positive correlation with F-actin levels. Our findings suggest that physical parameters of the microgels can be tuned to generate desired inflammatory response by modulating cytoskeletal dynamics of macrophages.

## 5. Summary and Conclusion

Our microgel system provides a controlled environment to study how physical characteristics can affect the phenotypes of adherent cells. The properties of the microgels can be easily tailored to evaluate a wide range of material properties, both independently and simultaneously. Evaluating the effects of curvature on macrophage phenotype provided insights into the influence of F-actin levels on the production of inflammatory mediators. This knowledge could inform the design of future biomaterials, enhancing therapeutic potential through the shape of the material alone. Additionally, materials can be designed to harness physical cues to modulate cell secretome and minimize the off-target effects of chemical mediators. Overall, our study highlights the importance of structural and material properties in biomaterial design and indicates that these properties can be leveraged for therapeutic use.

## 6. Acknowledgments

Research reported in this publication was supported in part by the National Institute of Arthritis and Musculoskeletal and Skin Diseases (NIAMS) Award Number R21AR078447, National Institute of General Medical Sciences (NIGMS) of the National Institutes of Health under Award Numbers P20GM130456, and Orthopedic Trauma Association (OTA, Grant Number: 6889). The content is solely the responsibility of the authors and does not necessarily represent the official views of the National Institutes of Health or other grant funding agencies.

